# Coupling Complementary Pathways in a Bacterial Co-culture Enables 4-Aminophenol Removal during Paracetamol Degradation

**DOI:** 10.64898/2026.11.11.765863

**Authors:** Francesca Demaria, Matthias Schaffter, Cristina Zimmardi, Marcel Suleiman, Pilar Junier, Philippe Corvini

## Abstract

Pharmaceutical biodegradation can generate potentially toxic metabolites that persist after the parent compound has been removed, posing additional risks to aquatic environments. Microbial co-cultures offer a promising strategy to address this challenge by combining complementary metabolic capabilities to remove both pharmaceuticals and their harmful transformation products by coupling complementary degradation pathways. Here, we investigated a two-species co-culture combining the paracetamol-degrading bacterium Delftia P1 with Microbacterium lacus to enable subsequent 4-AP removal. While Delftia P1 efficiently degraded paracetamol, it accumulated 4-AP as a dead-end metabolite. M. lacus was selected to complement this metabolic limitation because its genome contains a homolog of sadB, a monooxygenase gene involved in 4-AP transformation. The relative abundance of the two strains strongly influenced 4-AP removal, with the highest efficiency achieved at a Delftia P1:M. lacus cell-density ratio of 1:100, corresponding to an OD_600nm_ ratio of 1:10. At this ratio, the co-culture achieved complete removal of paracetamol and 4-AP within 6 and 24 h, respectively.

Its capacity was tested in wastewater, where 100% of paracetamol were removed within 4 h and no accumulation of 4-AP was observed within 24 h. These findings demonstrate the potential of cocultivating bacteria with complementary metabolic capabilities to address both pharmaceutical contamination and transformation-product accumulation. Moreover, the highest 4-AP removal was achieved when M. lacus was present at a 100-fold higher cell density than Delftia P1, highlighting the importance of considering relative cell densities when designing co-cultures.

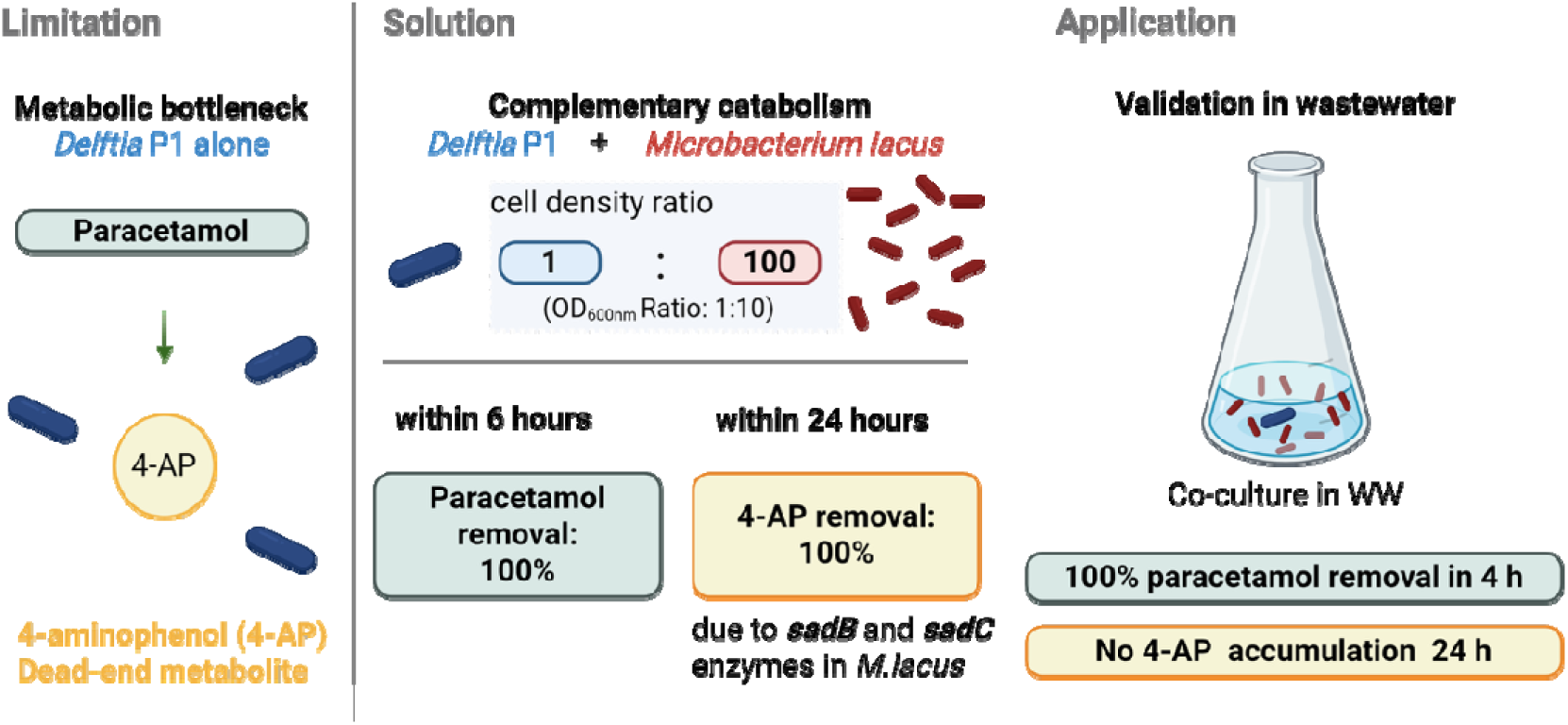

## 1. INTRODUCTION

Paracetamol, also known as acetaminophen, is one of the most widely consumed analgesic and antipyretic drugs worldwide (Albarano et al., 2025a; S. Wu et al., 2012; Yisau et al., 2024a). Its high consumption rates and continuous release from domestic, healthcare, and industrial sources have led to its widespread occurrence in the environment (Demaria et al., 2025; Gómez et al., 2006). As a result, paracetamol is now ubiquitously detected at ng/L to mg/L concentration ranges across diverse water environments, including wastewater treatment plant influents and effluents, hospital discharges, surface waters, groundwater, and occasionally drinking-water sources (Albarano et al., 2025b; Demaria et al., 2025; Hawash et al., 2023; S. Wu et al., 2012; Yin et al., 2025; Żur, Piński, et al., 2018). Although paracetamol generally shows relatively low acute toxicity at concentrations commonly detected in the environment, its persistence, high solubility, and continuous released, make its presence environmentally relevant (Żur, Wojcieszyńska, et al., 2018).

Moreover, pharmaceutical consumption is expected to increase in the coming decades, driven in part by population growth and, particularly, population aging, which is associated with a greater prevalence of chronic diseases and a higher demand for medication (Silva et al., 2021). Recent analyses estimate that the pharmaceutical market will grow at an average annual rate of approximately 6-8% through 2034 (Semchenkova, 2025). This increasing consumption is expected to translate into greater pharmaceutical loads in urban wastewater and, consequently, into aquatic environments (Silva et al., 2021). For example, paracetamol concentration increased 311 times in wastewater in Lanzhou (China) during the COVID-19 outbreak in comparison to paracetamol concentration pre-outbreak (Yu et al., 2025). Although this represents an exceptional period of increased consumption, it demonstrates how rapidly changes in pharmaceutical use can translate into substantially higher environmental concentrations.

These concerns apply not only for the parent compound, but also for their transformation products. Paracetamol degradation commonly begins with hydrolysis of the amide bond under aerobic conditions, leading to the formation of acetate, a readily assimilable carbon source, and 4-aminophenol (4-AP), a transformation product that can accumulate in the environment. Whenever possible, 4-AP is subsequently deaminated to hydroquinone; which undergoes further hydroxylation to form hydroxyquinol (1,2,4-trihydroxybenzene) (Lara-Moreno et al., 2024; Ricken et al., 2017). Hydroxyquinol is then subjected to dioxygenase-mediated aromatic ring cleavage, generating a linear intermediate (trans-2-hexenoic acid), that enters the TCA cycle (Lara-Moreno et al., 2024) (Fig. 1).

**Figure 1.**
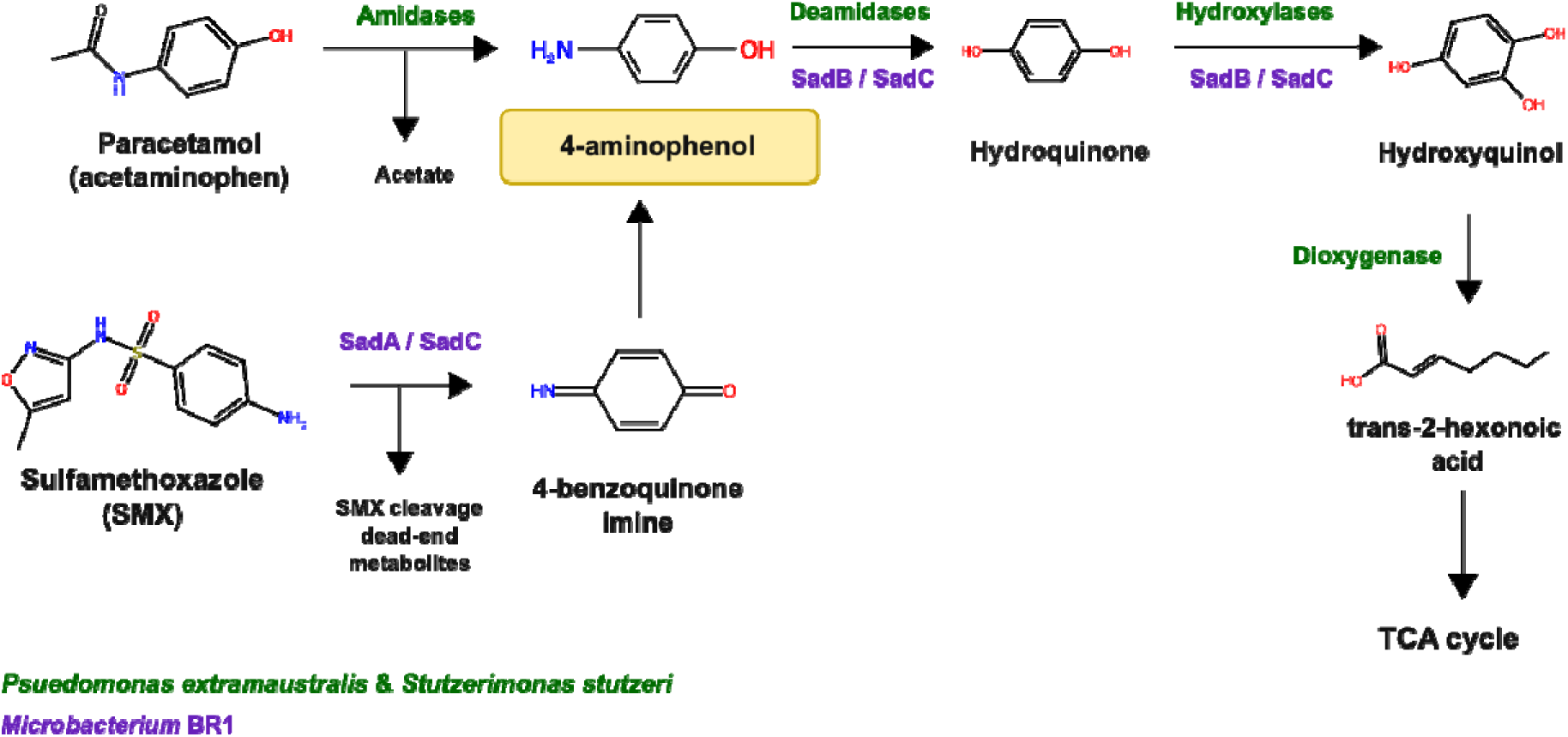
Degradation pathways of paracetamol in green (Lara-Moreno et al., 2024) and SMX in violet (Ricken et al., 2017) based on literature. Lara-Moreno *et al*. (Lara-Moreno et al., 2024) constructed a hypothetical paracetamol degradation pathway; based on genome sequencing and *in silico* prediction. According to this, in the first step, paracetamol gets converted to 4-AP through a hypothetical amidase-mediated cleavage of the acetamide bond. 4-AP is metabolized further down into hydroquinone, involving deamidases, and hydroquinol, involving hydroxylases, reaching the TCA cycle through trans-2-hexenoic acid due to the action of dioxygenases. The genetic basis of the SMX degradation pathway in *Microbacterium* BR1 was elucidated by Ricken *et al*. The *in vivo* analysis identified the *sad* gene cluster, which encodes two monooxygenases, SadA and SadB, together with the FMN reductase SadC. SadA and SadC catalyze the initial transformation of SMX, resulting in the release of 4-AP. This intermediate is then converted to 1,2,4-trihydroxybenzene (hydroxyquinol) by the combined activity of SadB and SadC prior to further mineralization (Ricken et al., 2017).

Among these intermediates, 4-AP has gained particular attention since recent evidence supports its accumulation in the environment. 4-AP was detected at µg/L concentrations across 20 wastewater treatment plants in China (Yin et al., 2025). Although it remains considerably less studied than paracetamol, 4-AP has raised concerns due to its potential toxicity and reactivity (Fdez-Sanromán et al., 2025; S. Wu et al., 2012; Żur, Wojcieszyńska, et al., 2018). It belongs to the broader class of phenolic compounds, which are known to interfere with enzymatic function and cellular metabolism, disrupt oxidative phosphorylation, alter antioxidant defense systems, and induce cellular damage (Ahmaruzzaman et al., 2024). These effects highlight the urge for effective degradation strategies that not only remove paracetamol from contaminated waters but also prevent the formation and accumulation of potentially harmful transformation products such as 4-AP.

Microorganisms can transform or mineralize pharmaceuticals, therefore microbial bioremediation is a promising and environmentally friendly approach (Demaria et al., 2025; Suleiman et al., 2023; Yisau et al., 2024b). The biodegradation of paracetamol has been reported in several bacterial isolates (Chen et al., 2022; Chopra et al., 2020; Poddar et al., 2022; Zhang et al., 2013b). For example, *Stenotrophomonas* sp. f1 and *Pseudomonas* strains fg-2 and f2 completely degraded concentrations ranging between 0.4 and 2.5 g/L (Zhang et al., 2013a). Other notable degraders include *Bacillus drentensis* S1, which removed 94.5 % of 0.3 g/L paracetamol within 48 h (Chopra et al., 2020), and *Pseudomonas* sp. PrS10, which achieved 96.37% degradation of 3 g/L within 168 h (Poddar et al., 2022). Similarly, *Shinella* sp. HZA2, completely degraded 0.1 g/L paracetamol within 12 h (Chen et al., 2022). Although individual bacterial strains efficiently degrade paracetamol, incomplete degradation pathways can limit full mineralization and promoting releasing 4-AP in the media.

Alongside paracetamol removal, Zhang et al. (2013a) and Chen et al. (2022) identified 4-aminophenol (4-AP) and hydroquinone as degradation intermediates, consistent with the reported catabolic pathways (Lara-Moreno et al., 2024). Identifying these intermediates does not establish their concentrations at the treatment endpoint, and paracetamol removal alone cannot demonstrate complete 4-AP elimination or mineralization. Poddar et al. (2022) reported no detectable 4-AP in an ethyl-acetate extract of the culture supernatant, but they did not report 4-AP-specific data or detection limits. Chopra et al. (2020) did not report 4-AP among the identified metabolites and observed brown-to-black coloration without identifying the compounds responsible. Under aerobic conditions, 4-AP can form insoluble brown material attributed to oxidative polymerization (Godain et al., 2020). However, coloration alone does not confirm this mechanism in the cultures discussed above, and further investigation is needed to distinguish biological degradation of 4-AP from abiotic transformation.

A possibility to maximize complete degradation is the use of several species with complementary capabilities. This is supported by metagenomic analyses performed on a bioreactor community indicating that different taxa contributed to full degradation by complementary catabolic functions. The initial transformations of paracetamol were performed by *Betaproteobacteria* and downstream ring-cleavage reactions performed by distinct community members, among which *Bacteroidetes, Rubrivivax, and Myxococcales_2* (Rios-Miguel et al., 2022). The enhanced biodegradation potential observed in natural microbial communities highlights the advantages of complementary metabolic functions, providing a rationale for the design of synthetic microbial communities that combine selected strains to achieve more efficient and complete biodegradation of the target substance. A successful example of this approach is a four-strain synthetic consortium that demonstrates efficient tetrabromobisphenol A (TBBPA) degradation through cooperative metabolism (T. Wu et al., 2025). Such interactions could prevent the accumulation of toxic intermediates and promote the complete degradation of target pollutants. Similarly, a consortium composed of one *Stenotrophomonas* and two *Pseudomonas* strains showed improved paracetamol degradation compared with the individual strains. At an initial paracetamol concentration of 2 g/L, the consortium achieved complete removal of the parent compound within 37 h and 87.1 % of the total organic carbon removal. In comparison, the individual *Pseudomonas* strains required approximately 45-60 h for complete paracetamol removal and achieved 48.9-65.2 % of the total organic carbon removal (Zhang et al., 2013a). These results highlight that the bacterial consortia could enhance the rate of pharmaceutical degradation and the removal of organic carbon from the culture medium. These examples have stimulated growing interest in bacterial co-culture to overcome the limitations of individual strains by combining the complementary degradation capacities of different species (Ruan et al., 2024).

Based on this rationale, we investigated the complementarity of bacteria with metabolic capabilities that could prevent the accumulation of 4-AP during paracetamol degradation. *Microbacterium* is a promising genus for the conversion of 4-AP to hydroquinone, as several members of this genus harbor key monooxygenase (Sad B) involved in 4-AP transformation (Li et al., 2025a; Ricken et al., 2017). *Microbacterium* BR1, was isolated for its capacity to degrade sulfamethoxazole (SMX). SMX leads to the release of 4-AP via ipso-substitution (Ricken et al., 2017). In *Microbacterium* BR1, the flavin-dependent monooxygenase SadB catalyzes further the oxidation of 4-AP to hydroxyquinol (trihydroxybenzene) (Fig. 1). Comparative genome analysis revealed the presence of a homologous *sadB* gene in the genome of *Microbacterium lacus*, suggesting a similar capacity to transform 4-AP (Li et al., 2025a).

In this study we aimed to develop a co-culture capable of degrading paracetamol with subsequent removal of 4-AP. The co-culture was based on *Delftia* P1 and *M. lacus. Delftia* P1 was isolated from a membrane bioreactor community adapted to a mixture of pharmaceuticals, then enriched in flasks on paracetamol only (Demaria et al., 2026; Suleiman et al., 2023). This strain degraded paracetamol efficiently but accumulated 4-AP as a dead-end metabolite. To overcome this limitation, *M. lacus*, which contained *sadB* in its genome, was selected (Li et al., 2025a; Tappe et al., 2013). We therefore hypothesized that co-cultivation of *Delftia* P1 with *M. lacus* would enable the complete removal of paracetamol and 4-AP from the culture. We hypothesized that *Delftia* P1 would convert paracetamol into 4-AP, while assimilating acetate, and *M. lacus* subsequently would transform 4-AP into hydroquinone thanks to the action of SadB. Furthermore, we proved that the degradation performance of 4-AP was strongly influenced by the cell ratio of *Delftia* P1 and *M. lacus*, with the highest 4-AP removal efficiency achieved at a 1:100 cell density *Delftia* P1:*M.lacus* ratio corresponding to a OD_600nm_ 1:10.

## 1. Materials and methods

### 1.1 Enrichment and Isolation of bacterial paracetamol degraders

A bacterial enrichment culture capable of degrading paracetamol was established from a microbial community previously adapted into a membrane bioreactor (MBR) to degrade a mixture of pharmaceuticals (atenolol, caffeine, diclofenac, enalapril, ibuprofen, and paracetamol each at a concentration of 100 mg/L) in a laboratory-scale membrane bioreactor (MBR) operated as described by Demaria *et al*. (Demaria et al., 2026).

Following adaptation, the microbial community was transferred to fresh synthetic wastewater (0.08 g/L peptone, 0.05 g/L meat extract, 0.015 g/L urea, 0.0035 g/L NaCl, 0.002 g/L CaCl□·2H□O, 0.0001 g/L MgSO□·7H□O, and 0.0014 g/L K□HPO□) containing paracetamol as the sole pharmaceutical at a concentration of 100 mg/L (662 µM). Batch enrichment cultures were initiated by inoculating 1% (v/v) of the adapted community into fresh medium and incubated for 2 days at 25°C with shaking at 160 rpm. At this point, paracetamol was fully removed. This enrichment procedure was repeated three successive times by transferring 1% (v/v) of each culture into fresh medium to promote the selection of specialized paracetamol-degrading microorganisms. After the third enrichment, serial ten-fold dilutions of the enrichment culture were prepared in sterile physiological saline solution (0.9 % NaCl), and 100 µL of the 10□□ and 10□□ dilutions were spread onto LB agar plates supplemented with 662 µM paracetamol. Plates were incubated at 25 °C overnight. Individual colonies displaying distinct morphologies were picked and purified by two successive rounds of re-streaking on fresh LB agar containing 662 µM paracetamol. All isolated strains were initially screened for their ability to degrade paracetamol (data not shown). Based on the results of this preliminary screening, the isolate P1 was selected for further experiments.

### 1.2 Bacterial identification

Paracetamol-degrading isolates were identified by colony PCR amplification using the universal bacterial primers 27F (5′-AGAGTTTGATCMTGGCTCAG-3′) and 1492R (5′-TACGGYTACCTTGTTACGACTT-3′), which amplify a 1200 bp approx. fragment of the bacterial 16S rRNA gene (Frank et al., 2008). Colony PCR was performed using the GoTaq® DNA Polymerase (Promega, Madison, WI, USA) in a final reaction volume of 50 µL containing 1× GoTaq buffer, 0.2 µM of forward and reverse primers, 0.2 mM of each dNTP (Solis Biodyne), 2.5 U of GoTaq DNA Polymerase, 2 µL of bacterial colony suspension, and nuclease-free water to the final volume. PCR amplification was carried out with an initial denaturation at 95 °C for 5 min, followed by 35 cycles of 95 °C for 30 s, 58 °C for 30 s, and 72 °C for 1 min 40 s, with a final extension at 72 °C for 5 min. PCR amplification and amplicon size were verified by agarose gel electrophoresis. Successfully amplified PCR products were purified using the QIAquick PCR Purification Kit (Qiagen, Hilden, Germany) according to the manufacturer’s instructions, and DNA concentrations were measured using a NanoDrop spectrophotometer (Thermo Fisher Scientific, Waltham, MA, USA). For sequencing, 12 µL of purified PCR product and 3 µL of the 27F primer (10 µM) were sent to Microsynth AG (Balgach, Switzerland) according to the manufacturer’s premix submission protocol. Amplicons were sequenced by Sanger sequencing, yielding approximately 1200 bp of the 16S rRNA gene. The resulting sequences were compared against the NCBI nucleotide database using BLASTn, and isolates were assigned to the closest matching taxon based on a minimum sequence identity of 99 % and a query coverage of 100%. Based on the taxonomic identification and institutional biosafety guidelines, only isolate P1, belonging to *Delftia* spp. with a Biosafety Level 1 taxa was retained for further experiments.

### 1.3 Paracetamol removal by the Delftia P1

Batch degradation experiments were performed in triplicate in a 50 mL Erlenmeyer flask at final working volume of 15 mL. The media used was synthetic wastewater (Section 1.1) supplemented with 630 µM paracetamol. *Delftia* P1 was grown overnight in 5 mL LB medium at 30 °C with shaking at 160 rpm, reaching a dilution-corrected optical density (OD_600nm_) of 2.58. Cells were harvested by centrifugation at 4000 × *g* for 10 min, washed once with sterile synthetic wastewater, and resuspended in synthetic wastewater supplemented with paracetamol prior to inoculation. Experimental flasks were inoculated with 580 µL of the washed preculture to reach an initial OD_600nm_ of 0.1. Cultures were incubated aerobically at 30 °C with shaking at 160 rpm. 500 µL of sample for quantification of paracetamol (Section 1.6), 125 µL of sample for 4-AP (Section 1.7), and 500 µL sample for flow cytometry analysis (Section 1.8) were collected from each flask every hour from 0 to 6 h, with an additional sample collected after 24 h. Microbial growth was monitored by measuring the optical density at 600 nm (OD_600nm_) of 250 µL samples using a BioTek Synergy S1 Multimode Reader. In parallel, bacterial cell abundance was quantified by flow cytometry using 500 µL samples and a CytoFLEX flow cytometer (Beckman Coulter, Brea, CA, USA), following the protocol described in Section 1.8. Control cultures were prepared under the same conditions in synthetic wastewater without paracetamol and inoculated at the same initial OD_600nm_. These controls were used to compare bacterial growth in the presence and absence of paracetamol.

### 1.4 Paracetamol and 4-AP removal by the co-culture in several cultivation conditions

Co-culture experiments were conducted to evaluate the effects of the initial *M. lacus* cell density and the presence or absence of SMX on paracetamol removal, 4-AP formation and removal. The co-culture consisted of *Defltia* P1 and *Microbacterium lacus* strain SDZm4 (DSM 18910) obtained from the German Collection of Microorganisms and Cell Cultures (DSMZ, Braunschweig, Germany) (National Center for Biotechnology Information, 2017). Upon receipt, the strain was revived according to the supplier’s instructions using DSMZ medium 514, incubated at 28 °C for 4 days, and stored at −80 °C in the corresponding growth medium supplemented with 30% (v/v) glycerol as a cryoprotectant.

To prepare the co-culture experiment, a preculture of *Delftia* P1. was prepared by inoculating 5 mL LB medium from a glycerol stock containing *Delftia* P1 and incubating overnight at 30 °C with shaking at 160 rpm. The preculture reached a dilution-adjusted optical density OD_600nm_ of 2.8. Prior to inoculation, the culture was harvested by centrifugation at 4000 g for 10 min, washed once with sterile synthetic wastewater, and resuspended in synthetic wastewater (Section 1.1) with SMX at 100 mg/L (395 µM) and paracetamol at 100 mg/L (662 µM) or only paracetamol at 662 µM, according to the experimental conditions.

*M. lacus* precultures were prepared in two separate bottles, each containing 100 mL of LB medium supplemented with 395 µM SMX. Cultures were incubated for 60 h at 25 °C, shaking at 160 rpm. The two precultures were subsequently pooled, resulting in a total culture volume of 200 mL with an OD_600nm_ of 0.70. The pooled culture was harvested by centrifugation at 4000 x g for 10 min washed once with sterile synthetic wastewater (Section 1.1), and concentrated by resuspension in a final volume of 45 mL of synthetic wastewater containing the compounds corresponding to the respective experimental conditions (SMX at 395 µM and paracetamol at 662 µM or only paracetamol at 662 µM). This concentration step increased the OD_600nm_ to approximately 3.0.

Co-culture paracetamol degradation experiments were performed in triplicates in 50 mL Erlenmeyer flasks containing final working volume of 15 mL. Two cultivation conditions were tested: paracetamol (662 µM) with SMX (395 µM) and paracetamol alone (662 µM). For each flask, 535 µL of the *Delftia* P1 preculture was added to obtain an initial OD_600nm_ of 0.1. Volumes of 0.5, 2.5, or 5.0 mL of the concentrated *M. lacus* suspension were added to obtain initial OD_600nm_ values of 0.1, 0.5, or 1.0, respectively. The final volume was adjusted to 15 mL with the corresponding synthetic wastewater medium (Section 1.1). Cultures were incubated aerobically at 30 °C with shaking at 160 rpm for 24 h.

Samples were collected at 2, 4, 6, and 24 h after inoculation. Aliquots of 500 µL were used for quantification of paracetamol and SMX by HPLC (Section 1.5), while 125 µL aliquots were used for spectrophotometric quantification of 4-AP (Section 1.6).

### 1.5 Degradation test by the co-culture in real wastewater

Co-culture degradation experiments were performed in triplicate using 50 mL Erlenmeyer flasks containing a final working volume of 15 mL of biologically treated wastewater (WW) supplemented with paracetamol as the sole pharmaceutical at an initial concentration of 630 µM. Wastewater was collected from the outlet of the moving bed bioreactor (MBBR) of the in-house wastewater treatment plant at the University of Applied Sciences and Arts Northwestern Switzerland (FHNW). Prior to the experiments, the wastewater was characterized by measuring pH, conductivity, chemical oxygen demand (COD), total organic carbon (TOC), total suspended solids (TSS), total nitrogen, and total phosphorus. The physicochemical characteristics of the wastewater are summarized in Supplementary Material **Error! Reference source not found.**.

*Delftia* P1 was cultivated overnight in LB medium, reaching a dilution-corrected OD_600nm_ of 1.745. *M. lacus* was cultivated in 100 mL of LB medium supplemented with SMX at 395 µM for 120 h, reaching a dilution-corrected OD_600nm_ of 2.65. Both cultures were harvested by centrifugation at 4,000 × g for 10 min, washed once with wastewater, and resuspended in wastewater supplemented with 600 µM paracetamol.

The co-culture was established at an initial *Delftia* P1:*M. lacus* OD_600nm_ ratio of 1:10. Specifically, 860 µL of the *Delftia* P1 suspension was inoculated to obtain an initial OD□□□ of 0.1, followed by 5.6 mL of the *M. lacus* suspension to obtain an initial OD_600nm_ of 1.0. The final volume was adjusted to 15 mL with wastewater supplemented with paracetamol. Cultures were incubated aerobically at 30 °C with shaking at 160 rpm. Samples were collected at 2, 4, 6, and 24 h after inoculation.

To assess sustained co-culture activity, paracetamol re-spiking experiments were performed in triplicate in 50 mL Erlenmeyer flasks at a final working volume of 20 mL. *Delfita* P1 and *M. lacus* were cultivated separately in LB medium overnight and in LB medium supplemented with 395 µM SMX for 120 h, respectively. Cells were harvested by centrifugation at 4,000 × *g* for 10 min, washed once with wastewater, and used to establish an initial *Delftia* P1:*M. lacus* OD_600nm_ ratio of 1:10. For *Delftia* P1, 1.7 mL of a preculture (having starting dilution-corrected OD_600nm_ of 1.13) was added to obtain an initial OD_600nm_ of 0.1. For *M. lacus*, 5.6 mL of a preculture (having a dilution-corrected OD_600nm_ of 3.51) was added to obtain an initial OD_600nm_ of 1.0. The cultures were incubated aerobically at 30 °C with shaking at 160 rpm. Paracetamol was re-spiked to a final concentration of 600 µM, with spikes occurring at 0, 6, 24, 30, and 48 h. The experiment was continued for 56 h. Samples were collected immediately before and after each re-spiking event and at intermediate sampling points for analysis of paracetamol degradation and 4-AP formation. Changes in the abundances of *Delftia* P1 and *M. lacus* were monitored by flow cytometry according to the procedure described in Section 1.8.

For the wastewater experiments, 500 µL aliquots were used for paracetamol quantification by HPLC (Section 1.5), 125 µL aliquots for spectrophotometric quantification of 4-AP (Section 1.6), and 500 µL aliquots for cell quantification by flow cytometry (Section 1.8).

### 1.6 HPLC analyses

Paracetamol and sulfamethoxazole (SMX) were quantified by high-performance liquid chromatography coupled with a diode array detector (HPLC-DAD). Prior to HPLC analysis, 500 µL were mixed with 125 µL of methanol and incubated at 4 °C for 20 min to precipitate proteins and maintain the analytes in solution. Samples were subsequently centrifuged at 21300 x g for 10 min at 4 °C, and the resulting supernatants were transferred to HPLC vials for analysis.

Chromatographic analyses were performed using an HPLC system (Agilent Technologies, Santa Clara, CA, USA) equipped with a diode array detector (DAD). Separation was achieved using an Infinity Poroshell 120 EC-C18 analytical column (3.0 × 100 mm, 2.7 µm; Agilent Technologies, Santa Clara, CA, USA) fitted with a ZORBAX Eclipse Plus C18 guard column (5 µm, 4.6 × 12.5 mm). The mobile phase consisted of solvent A (water containing 0.1% (v/v) formic acid) and solvent B (methanol). The gradient elution program was as follows: 0.0 - 1.0 min, 80% A / 20% B; 1.0 - 6.0 min, linear gradient to 70% A / 30% B; 6.0 - 6.5 min, linear gradient to 50% A / 50% B; and 6.5 - 7.0 min, linear gradient to 5% A / 95% B, followed by a 5 min re-equilibration period to restore the initial mobile phase composition. The flow rate was maintained at 550 µL min□¹, and the column temperature was set to 40 °C. Detection was performed at 280 nm using the DAD.

Paracetamol and SMX were identified by comparison of their retention times with those of analytical standards and quantified using external calibration curves. The curves were prepared from acetaminophen from Sigma standards ranging from 661.6 µM to to 2.5 µM, and SMX from Sigma standard ranging from 395 µM and 1.5 µM. Under these chromatographic conditions, paracetamol eluted at approximately 1.6 min. The limit of detection (LOD) and limit of quantification (LOQ) were 5.2 µM and 10.3 µM respectively. Calibration curves exhibited excellent linearity over the tested concentration range (R² = 0.9999). SMX eluted at approximately 4.8 min, with an LOD and an LOQ of 1.5 µM and a calibration curve exhibiting an R² value of 0.9998. Method precision was evaluated by six replicate injections of a standard solution, resulting in a relative standard deviation (%RSD) of less than 2%.

### 1.7 Detection and quantification of 4-AP

4-AP was quantified by fluorescence spectroscopy using a BioTek Synergy S1 Multi-Mode Reader (Agilent Technologies, Santa Clara, CA, USA) operated with excitation and emission wavelengths of 300 nm and 375 nm, respectively.

A calibration curve was prepared in synthetic wastewater for 4-AP quantificatio. Calibration standard curve of 4-AP (Sigma-Aldrich) were prepared at concentrations ranging from 916 µM to 3.6 µM. The stock solution was prepared under oxygen-free conditions by purging with nitrogen to prevent early oxidation, polymerization, and precipitation of 4-AP, as previously reported (Rios-Miguel et al., 2022). Calibration standards were freshly prepared prior each analysis. At each sampling time point, 125 µL culture aliquots were collected and centrifuged at 21300 x g for 10 min at 4 °C to remove bacterial cells and suspended solids. The resulting supernatants were diluted 1:1 (v/v) with methanol according to the protocol described by *Dejaegher et al*. (Dejaegher et al., 2008). Fluorescence measurements were performed using a BioTek Synergy S1 Multi-Mode Reader (Agilent Technologies, Santa Clara, CA, USA) with excitation and emission wavelengths of 300 nm and 370 nm, respectively. The calibration curve exhibited a linear response over the tested concentration range, with a correlation coefficient (R²) of 0.9756. The limit of detection (LOD) and limit of quantification (LOQ) were 3.57 µM.

### 1.8 Flow cytometry

Flow cytometry was used to quantify the cell densities of *Delftia* sp. and *M. lacus* in the co-culture throughout the degradation experiments. Samples were collected from each biological replicate at 0, 2, 4, 6, and 24 h. Aliquots of 500 µL were centrifuged at 4000 × *g* for 10 min at 24°C, washed once with sterile physiological saline solution (0.9% NaCl), and resuspended in 500 µL of physiological saline to reduce background signal.

For method optimization, samples were analyzed undiluted as well as at serial ten-fold dilutions (10^-1^, 10^-2^, and 10^-3^). Based on these preliminary experiments, samples were routinely analyzed at a dilution of 10^-2^. Cells were stained with SYBR Green I (Invitrogen, Thermo Fisher Scientific, Waltham, MA, USA). The commercial stock solution 10000x was diluted to 25x to prepare the working solution, and 10 µL of working solution was added to 240 µL of each sample to obtain a final dye concentration of 1x. Samples were incubated for 10 min at 37 °C in the dark prior to analysis.

Flow cytometric analyses were performed using a CytoFLEX flow cytometer (Beckman Coulter, Brea, CA, USA) equipped with 405, 488, and 638 nm lasers. Data were acquired at a flow rate of 10 µL/min, and 2 µL of each sample was analyzed. Data acquisition and analysis were performed using CytExpert software (Beckman Coulter).

Bacterial populations were identified using a sequential gating strategy. An initial gate based on forward scatter (FSC) and side scatter (SSC) was used to discriminate bacterial cells from large particles and electronic noise. SYBR Green fluorescence was subsequently used to distinguish bacterial cells from non-cellular debris, particularly for *M. lacus*, whose scatter characteristics partially overlapped with those of background particles (Supplementary Material **Error! Reference source not found.**). An additional gate based on SYBR Green fluorescence versus side scatter height (SSC-H) was applied to further exclude background events.

The gating strategy was established using unstained samples, monocultures of *Delftia* sp. and *M. lacus*, and abiotic controls. To validate the gating strategy, the number of detected events per microliter at the inoculation density was compared with the corresponding colony-forming units (CFU/mL) (Supplement material **Error! Reference source not found.**) estimated from optical density measurements. Agreement between flow cytometry counts and CFU estimates was used to confirm that the gated events corresponded to bacterial cells. Moreover, the cell dimension was evaluate using SEM to understand better the OD_600nm_ discrepancies with the cell counts (Supplemntary material **Error! Reference source not found.**).

### 1.9 Statistical analysis

A two-way analysis of variance (ANOVA) was performed using GraphPad Prism version 10.4.1 (GraphPad Software, 2024) to evaluate the effects of paracetamol and incubation time on *Delftia* P1 growth. The statistical model included paracetamol treatment and incubation time as fixed factors, as well as their interaction. Statistical significance was defined as *p* < 0.05.

The effects of SMX presence or absence and three initial *M. lacus* inoculation densities (OD□□□ of 0.1, 0.5, and 1.0) on 4-AP removal in the co-cultures were statistically evaluated. Three replicates were analyzed for each experimental condition.

For each replica, the percentage of 4-AP removed at each sampling time was calculated as: (Supplementary Materials **Error! Reference source not found.**):

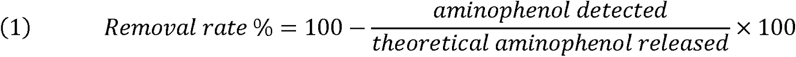

where the measured 4-AP concentration (µM) was determined experimentally by fluorescence analysis (Section 1.7), and the theoretical 4-AP concentration (µM) was calculated assuming complete stoichiometric conversion (1:1 molar ratio) of the paracetamol and/or SMX trasformed into 4-AP at each sampling point (Supplementary Material **Error! Reference source not found.**). Because 4-AP was detectable only during the first 6 h before precipitation occurred, the average removal rate (% /h) was calculated by dividing the removal percentage measured over this period by 6 h.

The effect of the initial *Microbacterium lacus* cell density on the 4-AP removal rate was evaluated using a one-way analysis of variance (ANOVA). Additionally, the combined effects of the initial *M. lacus* cell density and the presence or absence of sulfamethoxazole (SMX) on the 4-AP removal rate were assessed using a two-way analysis of variance (ANOVA), including the interaction between both factors. Statistical significance was defined as *p* < 0.05. Statistical analyses were performed using GraphPad Prism version 10.4.1 as described more in details in the section 1.9 (GraphPad Software, 2024).

## 2 Results

### 2.1 Paracetamol degradation by Delftia

Bacteria degrading paracetamol were isolated from membrane bioreactor communities running on synthetic wastewater containing a mixture of pharmaceuticals at concentration of 100 mg/L each. For this, the enrichment cultures were set with paracetamol as sole pharmaceutical. After four generations of enrichment, five isolates were isolated for their paracetamol capabilities and sequenced. Among them, Delftia P1 was selected for further analysis because it consistently showed the highest and fastest degradation capacity, removing paracetamol completely within 24 h at 662 and 3310 µM (data not shown).

The degradation of paracetamol by *Delftia* P1 was studied in detail by monitoring paracetamol concentration, 4-AP formation, and cell numbers over time (Fig. 2a, 2b). Paracetamol was fully degraded within 4 h, whereas 4-AP accumulated simultaneously at an equimolar concentration. This pattern is consistent with the literature, which reports that paracetamol degradation is initiated by an amidase-mediated hydrolysis, likely involving a hypothetical extracellular amidase (Abada, 2014), releasing 4-AP and acetate as the first degradation products (Fig. 2c) (Rios-Miguel et al., 2022). The acetate released during this reaction could support bacterial growth. Accordingly, in the cultures supplemented with 662 µM paracetamol, *Delftia* P1 showed a five-fold increase in cell density within 2 h, rising from around 1.4 × 10^7^ to 5 × 10^7^ cells, whereas no comparable increase was observed in cultures grown without paracetamol (control) (Fig. 2b). Two-way ANOVA revealed significant effects on *Delftia* P1 growth on paracetamol (*p* < 0.0001), time (*p*< 0.0001), and the interaction between time and paracetamol treatment (*p*< 0.0001). This indicates that *Delftia* growth over time increased significantly in the presence of paracetamol. After 24 h (data not shown), 4-AP levels decreased again presumably because of oxidation, polymerization, and accumulation as a brown precipitate (Rios-Miguel et al., 2022) (Fig. 2d). Together, these results demonstrate the high paracetamol-degrading efficiency of *Delftia* sp. while also revealing the transient accumulation of 4-AP during the degradation process (Fig. 2c).

**Figure 2.**
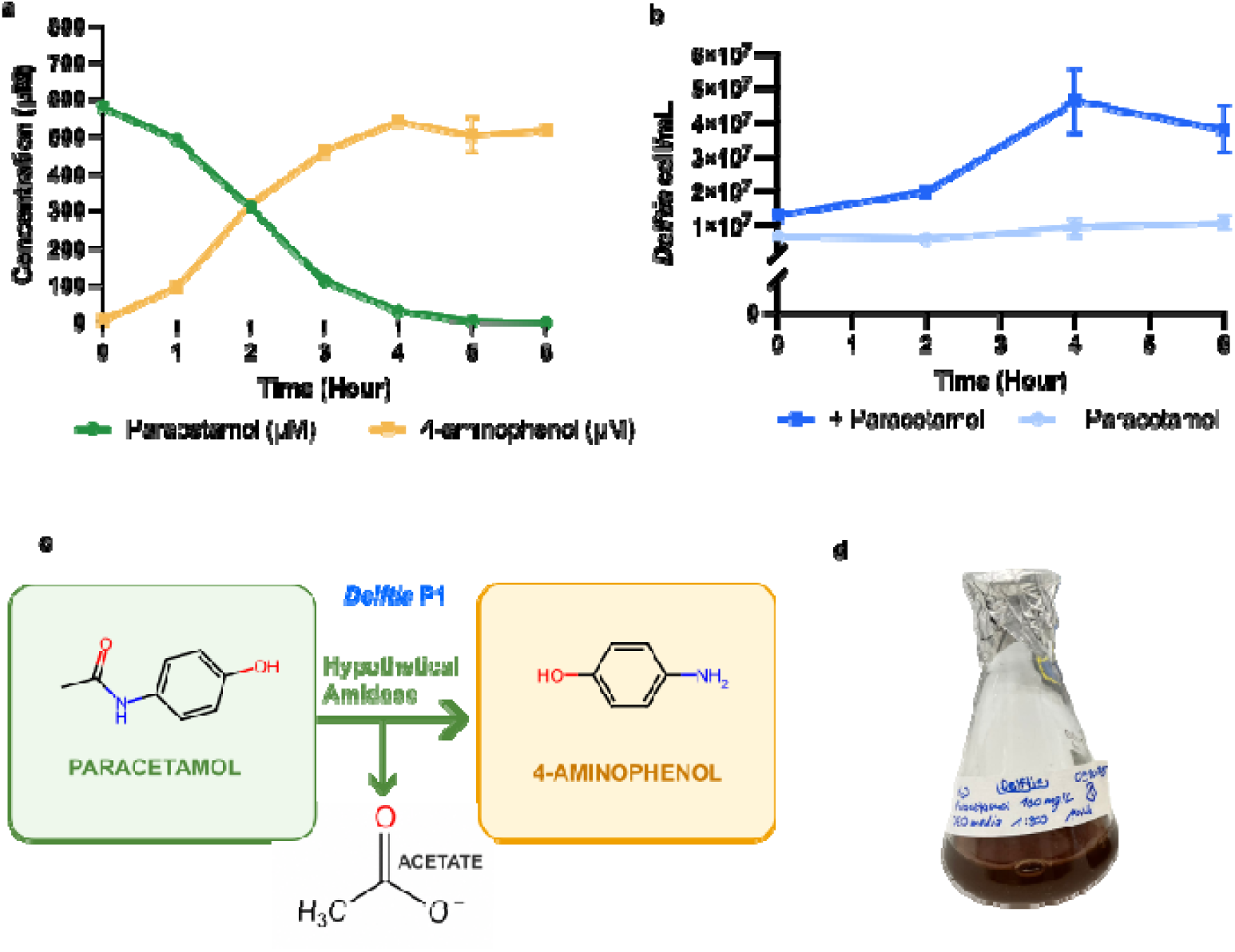
Partial degradation of paracetamol in the presence *Delftia* P1. **a** Degradation of paracetamol and accumulation of 4-AP over 6 hours. **b** *Delftia* P1 cell count (*Delftia* cell/mL) was assessed using flow cytometry. The cellular growth in the presence of paracetamol was measured over 6 hours (dark blue curve). A negative control with *Delftia* cultivated in absence of paracetamol was also included (light blue curve). **c** Schematic representation of the degradation pathway by *Delftia* P1. showing the precursor (paracetamol) and the metabolites (4-AP and acetate). **d** *Delftia* P1 culture showing a brown precipitate after 24 hours of incubation, presumably due to oxidation/polymerization of 4-AP.

### 2.2 Co-culture potential for paracetamol degradation

The combination of *Delftia* P1 and *M. lacus* in a co-culture was considered as a promising synergistic strategy for improving paracetamol degradation. This approach was motivated by the presence of the *sad* gene cluster in the genome of *Microbacterium* spp., particularly the presence of the *sadB* gene encoding a flavin-dependent monooxygenase involved in the transformation of 4-AP generated during SMX degradation (Li et al., 2025a; Ricken et al., 2015) (Fig. 3a). Genome analysis revealed that *M. lacus* strain SDZm4, which is also capable of degrading sulfonamides, harbors genes encoding flavin-dependent monooxygenases, making it a suitable candidate for investigating the role of these enzymes in 4-AP degradation (Li et al., 2025a). Since *Delftia* P1 rapidly converts paracetamol into 4-AP, whereas *M. lacus* was expected to further metabolize this intermediate, both strains were cultivated together to evaluate whether 4-AP accumulation could be reduced. Co-cultures were tested in synthetic wastewater containing either paracetamol and SMX or paracetamol alone. *Delftia* P1 sp. was inoculated to reach an OD_600nm_ of 0.1, while *M. lacus* was added to reach different initial cell densities (OD_600nm_ 0.1, 0.5, or 1.0). This setup was used to determine whether the presence of SMX and the initial abundance of *M. lacus* influenced paracetamol degradation and 4-AP accumulation.

**Figure 3.**
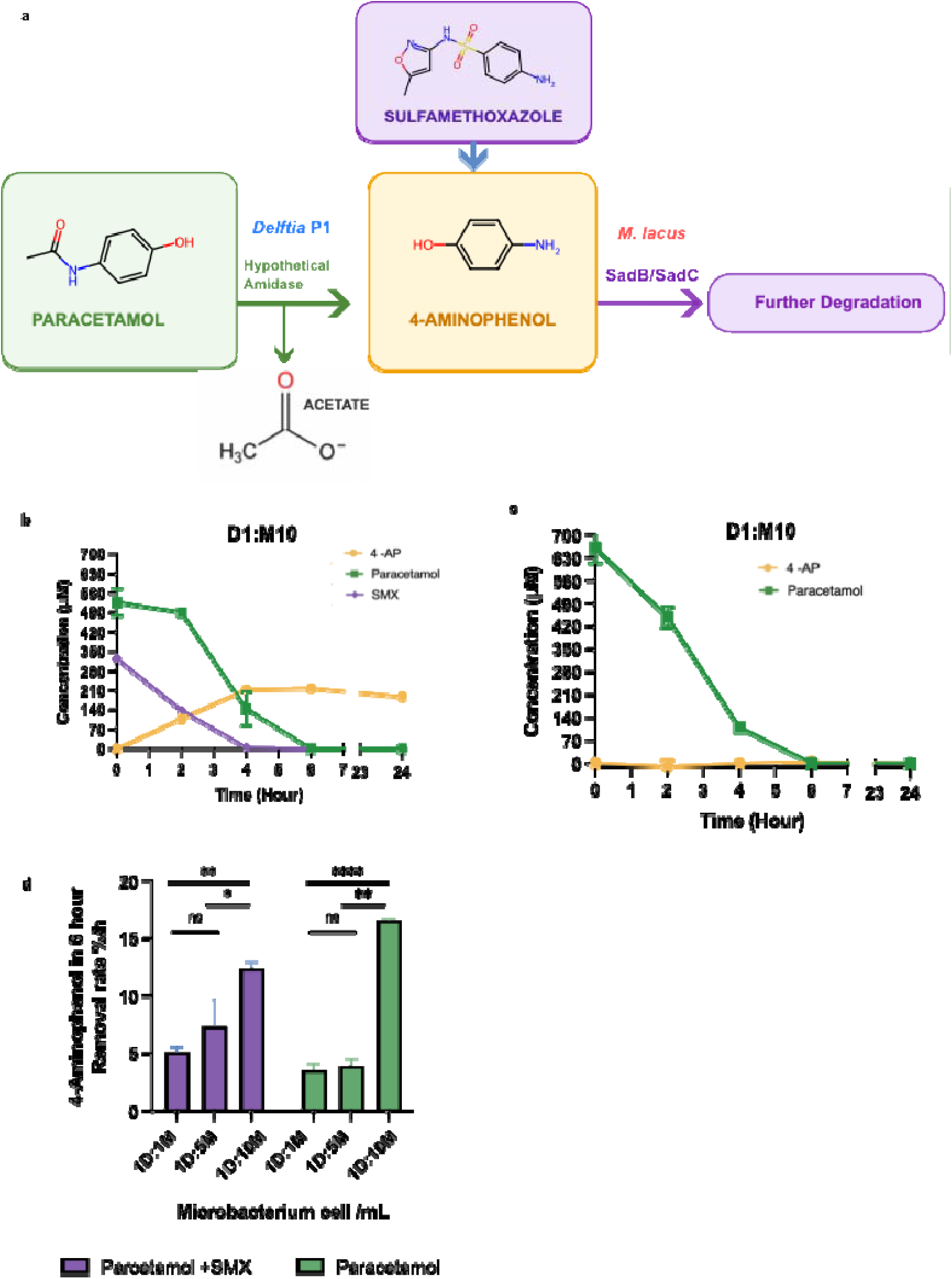
Removal of paracetamol and 4-AP by a co-culture of *Delftia* P1 (initial OD_600nm_ = 0.1) and *M. lacus* (initial OD_600nm_ = 1). **a** Schematic of the proposed synergistic interaction: *Delftia* P1 converts paracetamol (green) to 4-AP (yellow), while *M. lacus* further metabolizes 4-AP, presumably by the action of SadB. SMX degradation also contributes to 4-AP formation (violet). **b-c** Paracetamol and 4-AP removal by the co-culture at 1:10 *Delftia* P1: *M.lacus* OD ratio with (**b**) or without (**c**) SMX. Additional ratios are shown in Supplementary Materials **Error! Reference source not found.**. **d** 4-AP removal (%h^-1^) for the three tested ratios (D1:M1, D1:M5, D1:M10) in the presence or absence of SMX. Statistical differences were assessed by one-way ANOVA followed by Tukey’s multiple-comparison test and are indicated by *,**, and ***.

Paracetamol was completely hydrolyzed within 6 h under all experimental conditions (Fig. 3b; Supplementary Materials **Error! Reference source not found.**). However, cultures inoculated with the highest *M. lacus* cell density (OD_600nm_ 1) (Figure 3 b and c), exhibited a slight delay in paracetamol degradation, with residual concentrations of approximately 109–142 µM after 4 h, compared with only 20–40 µM in the other co-culture ratios (Supplementary Materials **Error! Reference source not found.**). These observations indicate that increasing the initial abundance of *M. lacus* slightly affected the kinetics of paracetamol degradation without altering the final removal efficiency. In contrast, SMX degradation strongly depended on *M. lacus* cell density, with higher inoculum densities resulting in faster and complete SMX removal (Supplementary Materials **Error! Reference source not found.**, Figure 6 b). A strong effect of the co-culture composition was observed concerning the concentration of the intermediate 4-AP. At low *M. lacus* abundances, 4-AP accumulated in the culture and progressively precipitated, forming brown precipitates. 4-AP removal reached 100% only when *M. lacus* was inoculated at a ten-fold higher OD_600nm_ than *Delftia* P1 in culture containing paracetamol alone (Fig. 3c), indicating that higher *M.lacus* cell density is required for efficient 4-AP removal.

To quantify these differences, the 4-AP removal rate (%/h) was compared among the three inoculum ratios of *M. lacus* (OD_600nm_ 0.1, 0.5, 1.0) using one-way ANOVA followed by Tukey’s multiple-comparisons test. Initial *M. lacus* abundance significantly affected the removal rate in both the paracetamol-only (*p* = 0.0016) and paracetamol + SMX (*p* < 0.0001) treatments. In both the presence and absence of SMX, Tukey’s test showed that cultures inoculated with the highest *M. lacus* cell density ratio had significantly higher 4-AP removal rates over the 6 h incubation period. At the highest starting cell density, the removal rate reached approximately 12%/h in SMX-supplemented cultures and 15%/h in cultures supplied with paracetamol only. In contrast, at the two lower starting inocula (OD_600nm_ = 0.1 and 0.5), removal rates were approximately 7 and 4.5%/h, respectively, in the SMX-supplemented cultures, and around 3–4%/h in the paracetamol-only cultures (Fig. 3d).

Because Sad B has been associated with sulfamethoxazole degradation, we further investigated whether SMX could enhance 4-AP removal. Preliminary experiments confirmed that SMX neither inhibited the growth of *Delftia* P1 nor paracetamol inhibit the growth of *M. lacus* (Supplementary Material **Error! Reference source not found.**). A two-way ANOVA, using *M. lacus* cell concentration and SMX presence as independent factors, revealed that *M. lacus* abundance was the primary determinant of 4-AP degradation (*p* < 0.0001), whereas SMX alone had no significant effect (*p* = 0.2901). A significant interaction between *M. lacus* starting cell density and SMX presence was observed (*p* = 0.0007), indicating that the influence of cell density differed slightly depending on the treatment condition (with or without SMX). Tukey’s post hoc analysis confirmed that the highest *M. lacus* inoculum ratio (OD_600nm_ = 1.0) achieved significantly greater 4-AP removal than the two lower inoculum levels under both conditions (around 12 -12 %/h vs 4 – 7 %/h), whereas no significant differences were observed between OD_600nm_ values of 0.1 and 0.5.

Although 4-AP removal reached 70% in the presence of SMX, the highest removal efficiency (100%) was observed in cultures containing paracetamol alone. This difference is likely explained by the fact that SMX, which degradation also produces 4-AP contribute to increasing the pool of 4-AP, limiting or slowing down the catabolism of the latter. Complete degradation of SMX occurred within 4 h, and this should theoretically produce approximately 59% more 4-AP than paracetamol alone, thereby increasing the intermediate load that must be metabolized by the co-culture. This can explain why a lower 4-AP removal was achieved in the presence of SMX, despite the higher *M. lacus* cell density.

Overall, these findings demonstrate that the initial abundance of *M. lacus* is the principal factor governing 4-AP removal, whereas SMX does not enhance the overall degradation efficiency. Based on these results, the subsequent wastewater experiments were performed using the co-culture ratio of *Delftia* P1:*M. lacus* (OD_600nm_ 0.1:1) without the addition of SMX.

### 2.3 Co-culture degrading capability and dynamics in real wastewater

We applied the co-culture to wastewater where paracetamol was added in order to assess its functionality in a real matrix. In a real biologically treated wastewater, the co-culture with *Delftia* P1 and *M. lacus* in a 1:10 OD_600nm_ ratio remained fully functional and efficiently removed paracetamol (Fig. 4a). Complete paracetamol degradation was achieved within 4 h, with the major decrease concentration occurring already after 2 h. The rapid decay of paracetamol (Fig. 4a, green curve) was accompanied by a transient increase in 4-AP, reaching 64 µM at hour 2 (Fig. 4a, yellow curve), followed by its subsequent decline, indicating effective biodegradation of this intermediate. No brown coloration was observed in the wastewater matrix, further suggesting that 4-AP did not accumulate in the medium. In the control, both paracetamol remained unchanged and 4-AP did not accumulate throughout the experiment (Fig. 4a, black and grey curves, respectively). Overall, these results demonstrate that the co-culture maintained its synergistic activity in real wastewater and effectively prevented the accumulation of the toxic intermediate 4-AP.

**Figure 4.**
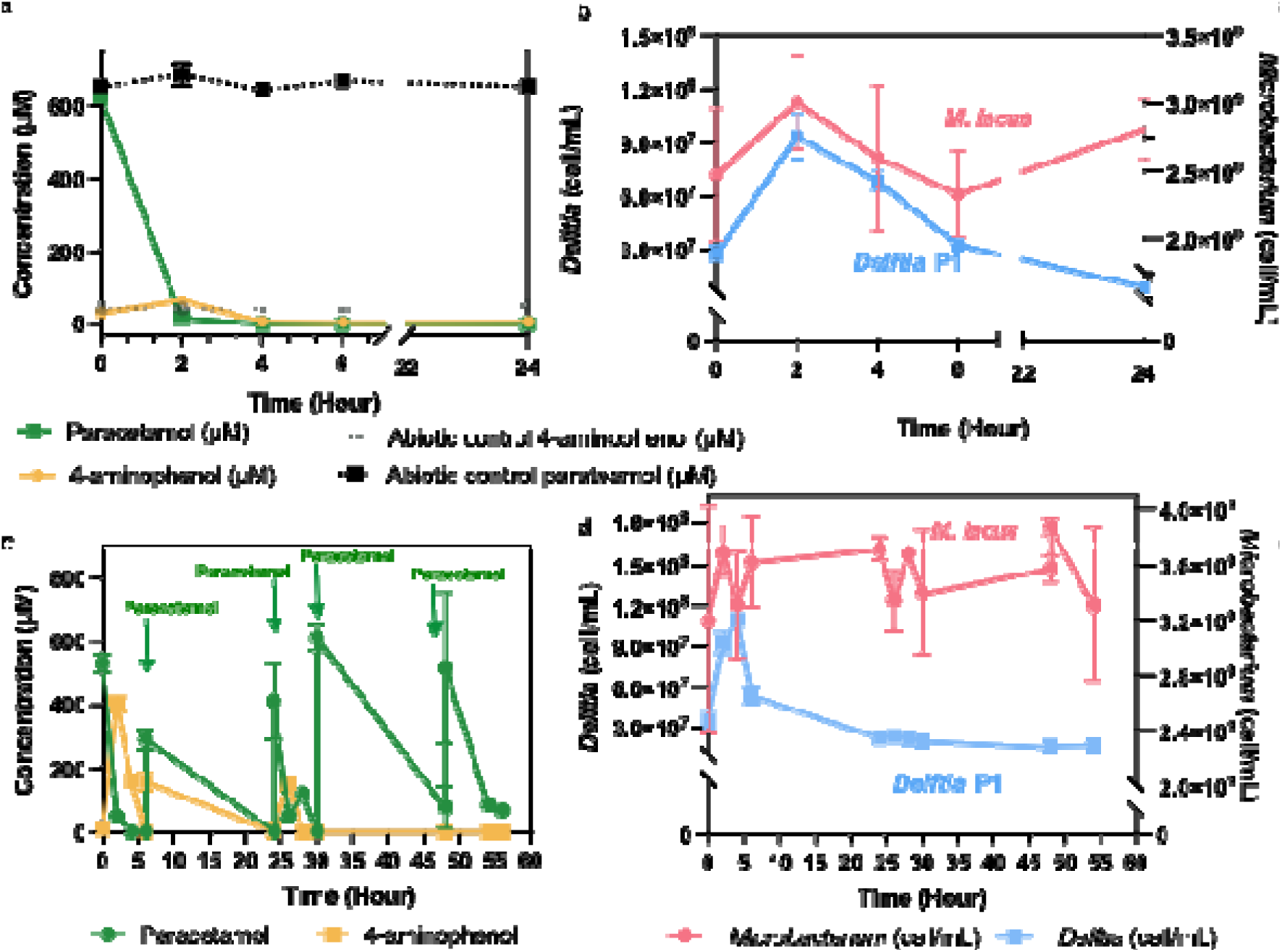
Paracetamol and 4-AP removal by the co-culture (*Delftia* P1 :*M. lacus,* 1:10) in real biologically treated wastewater over 24 and 56 hours. **a** Concentrations of paracetamol (green curve) and 4-AP (yellow curve) during treatment with the co-culture in biologically treated wastewater. The abiotic control is shown by the black (paracetamol) and grey (4-AP) curves. **b** Cell counts of both bacterial species were monitored over 24 h using flow cytometry. **c** Paracetamol degradation over time, after paracetamol re-spike of around 600 µM at 6, 24, 30, 48 hours. The green curve represents the paracetamol concentration; the yellow curve the 4-AP concentration over56 hours. The paracetamol removal (green curve) occurred at each paracetamol re-spike. **d** Bacterial cell count (cell/mL) was performed using flow cytometry *Microbacterium* represented by the red curve, *Delftia* P1 represented by the blue curve.

Flow cytometry data showed that the main phase of paracetamol removal, occurring after 2 h, coincided with an increase in *Delftia* P1 cell numbers (Fig. 4b, blue line). *Delftia* P1 increased approximately five-fold, from 2 x 10^7^ to 1 x 10^8^ cells within the first 2 h, before declining at later sampling times. This increase is consistent with the uptake of acetate as a carbon source during paracetamol degradation. In contrast, *M. lacus* cell numbers did not increase significantly over the 24 h cultivation period, probably due to its slow growth rate or high cell density.

Interestingly, the initial cell density of *M. lacus* was approximately 100-fold higher than that of *Delftia* P1 ratio, 10 times more than expected from the OD_600nm_. Further analysis correlating colony-forming units with OD_600nm_ values (Supplementary Materials **Error! Reference source not found.**) confirmed that, at the same optical density, *M. lacus* contained a higher number of cells than *Delftia* P1. This difference may be partially explained by the smaller cell size of *M. lacus* compared with *Delftia* P1 (*i.e.* approximately 0.5 µm and 1.6 µm respectively (Supplementary Material **Error! Reference source not found.**)).

To further evaluate the robustness and long-term functionality of the co-culture, we tested *Delftia* P1 and *M. lacus* under repeated paracetamol re-spiking conditions over three days. This experimental setup was considered to assess whether the co-culture could remove paracetamol and 4-AP over time after repeated substrate exposure. Although these conditions do not fully represent a realistic wastewater treatment scenario, where a continuous system would likely be more appropriate, the results showed a clear and reproducible degradation capacity (Fig 4c). After each paracetamol re-spiking event, the co-culture removed paracetamol within the first 2 h. For the overnight intervals, intermediate data points were not collected; however, samples taken the following morning confirmed that paracetamol had been degraded. The formation of 4-AP was observed during the first 2 h after re-spiking, reaching up to 400 µM, then the intermediate was subsequently removed. Interestingly, during later re-spiking cycles, 4-AP was detected at lower extent and was still efficiently degraded, suggesting that the co-culture maintained, and potentially improved, its ability to process the toxic intermediate over time. Regarding cell dynamics (Fig. 4d), *Delftia* P1 showed an initial increase in cell numbers within the first 2–4 h, corresponding to the first addition of paracetamol at inoculation time, before respiking. However, no further growth was detected during the following re-spiking cycles. This suggests that paracetamol degradation may have continued through persistent enzymatic activity, potentially due to amidase released into the supernatant, even when *Delftia* P1 cells were no longer actively growing or fully fit. In contrast, *M. lacus* did not show a consistent increase or decrease in cell density throughout the experiment, indicating a relatively stable population. Since propidium iodide staining was not used, we could not directly assess cell viability and fitness or determine whether part of the population was dying. Nevertheless, no substantial debris formation was observed, suggesting that extensive cell lysis was unlikely. Overall, these results indicate that the co-culture retained its functional stability under repeated paracetamol exposure and consistently prevented long-term accumulation of 4-AP in treated wastewater.

## Discussion

One of the major environmental challenges is the contamination of water sources by pharmaceuticals and their metabolites, which can persist through wastewater treatment and enter aquatic environments. One abundant pharmaceutical is paracetamol: while the parent compound is readily degraded to 4-aminophenol (4-AP), this intermediate can persist and accumulate, posing a challenge for the complete removal of this pollutant. Consistent with this concern, Yin et al., (2025) detected 4-AP in all 20 wastewater treatment plant samples they analyzed, with effluent concentrations exceeding those of paracetamol. Together, these findings point to an imbalance: wastewater communities may transform paracetamol faster than they remove the 4-AP produced. These findings indicate that the effective elimination of 4-AP is a more relevant measure of treatment performance than the disappearance of the parent compound alone. To address this challenge, this study investigated a co-culture combining a strain capable of rapid paracetamol degradation with a 4-AP-consuming strain to promote the removal of both the parent compound and the intermediate.

While *Delftia* P1 rapidly transformed paracetamol, consistent with previous reports for other strains (Rios-Miguel et al., 2022), this transformation led to the accumulation of 4-AP in monoculture. This illustrates a broader challenge in paracetamol degradation, while many bacteria can initiate the process, they produce toxic intermediates that they cannot further degrade (Rios-Miguel et al., 2022; Yin et al., 2025). A consortium can address this type of bottleneck by combining organisms with complementary metabolic capabilities to remove intermediates, recalcitrant to biodegradation (Demaria, et al., 2026; Lara-Moreno et al., 2024; T. Wu et al., 2025). The benefit of combining strains was also demonstrated by Zhang et al., (2013), whose consortium of *Stenotrophomonas* sp. f1, *Pseudomonas* sp. f2, and *Pseudomonas* sp. fg-2 outperformed the individual cultures, achieving paracetamol mineralization. However, the contributions of the individual strains to specific degradation steps remained unresolved. Our study provides this functional distinction: *Delftia* P1 rapidly transformed paracetamol, while *M. lacus* subsequently removed the resulting 4-AP, preventing the intermediate accumulation observed in the *Delftia* P1 monoculture. A key strength of our study is therefore the identification of complementary strain-specific roles in the removal of paracetamol and 4-AP. These findings suggest complementary interactions among the strains could be a valuable solution for pharmaceuticals removal (Lara-Moreno et al., 2024; Zhang et al., 2013b).

The rationale for this co-culture lies in repurposing the *sad* gene cluster of *Microbacterium*, originally characterized in sulfamethoxazole degradation, to remove 4-AP generated during paracetamol degradation. Within this pathway, SadB and SadC enable the transformation of 4-AP into 1,2,4-trihydroxybenzene (Ricken et al., 2017), providing a metabolic link between the degradation pathways of two different pharmaceuticals. Given the presence of genes encoding SadB and SadC in the *M. lacus* genome, we hypothesized that this strain could complement *Delftia* P1 by transforming the accumulated 4-AP. The improved 4-AP removal observed in co-culture was consistent with this hypothesis; however, whether 4-AP was completely metabolized remains unclear.

One key finding was that successful transformation of 4-AP in co-culture depended on the initial cell density ratio of the two strains, highlighting the importance of an appropriate inoculum composition for rapid and reliable removal. At the selected *Delftia* P1:*M. lacus* inoculation cell density ratio of 1:100, the removal of paracetamol without detectable 4-AP accumulation occurred. The consortium at the same cell density ratio maintained the paracetamol removal and downstream 4-AP removal observed in synthetic wastewater when applied to biologically treated wastewater, where no 4-AP accumulation was detected, suggesting that its complementary transformation capacity remained effective in a more complex matrix. Paracetamol hydrolysis releases 4-AP and acetate without requiring a cofactor, whereas SadB-mediated 4-AP transformation requires oxygen and reduced FMN supplied by a reductase such as SadC (Ricken et al., 2017). A higher initial abundance of *M. lacus* may therefore be required to provide sufficient catalytic capacity for efficient 4-AP removal. Similar effects about the role of the cell ratio have been observed in other studies. Jayachandran et al., (2009) reported that inoculum composition affected degradation kinetics and catechol accumulation during the simultaneous degradation of phenol and 3-chlorobenzoate by two *Pseudomonas* strains. Qi et al., (2021) likewise demonstrated the importance of inoculum composition in a consortium coupling PET depolymerization with downstream terephthalate consumption.

In co-culture, repeated paracetamol additions resulted in continued degradation without detectable 4-AP accumulation, although *Delftia* P1 showed no further growth after the first addition. We hypothesize that *Delftia* P1 metabolized the acetate released during paracetamol hydrolysis to support cellular maintenance without further net growth.

We also tested the addition of sulfamethoxazole (SMX) to the co-culture to test whether 4-AP removal could be improved at lower cell densities ratio. Sulfonamides are known to enhance *sadA* and *sadB* expression in *Microbacterium* sp. HA-8 (Li et al., 2025b). Because SadB participates in 4-AP transformation, this finding provided a rationale for testing SMX supplementation in our system. However, SMX addition did not affect 4-AP removal under any of the tested conditions.

Several factors may explain why SMX addition did not improve 4-AP removal. The induction of *sadAB* reported in *Microbacterium* sp. HA-8 may not occur to the same extent in *M. lacus*, and gene expression and enzyme activity were not measured to verify this response in our strain. Even if induction occurred, increased gene expression would not necessarily translate into faster 4-AP transformation. Furthermore, SMX degradation by *M. lacus* provides an additional source of 4-AP, increasing the total load requiring downstream transformation. This additional production could offset increased transformation capacity and mask a stimulatory effect of SMX in the measured 4-AP concentration profiles.

This study demonstrates that combining strains with complementary metabolic capabilities can improve the removal of a pharmaceutical and its degradation intermediate. Pairing *Delftia* P1 with *M. lacus* enabled rapid paracetamol transformation and subsequent 4-AP removal, with the initial cell density ratio emerging as a key factor for efficient degradation. Continued degradation following repeated paracetamol additions further supports the potential of this co-culture for sustained treatment. However, confirming complete mineralization and detoxification will require carbon balances, identification of remaining transformation products and toxicity measurements. Longer-term studies under continuous loading will be essential to assess whether these benefits translate into reliable performance in biological wastewater treatment.

## Conclusion

Taken together, the results support rational consortium assembly as a promising strategy for addressing pharmaceutical transformation bottlenecks in wastewater. The value of this approach is the deliberate pairing of a fast initial transformer, *Delftia* P1, with a partner selected for a plausible downstream 4-AP transformation capacity. The observed dependence on inoculation ratio further shows that consortium performance depends on matching pathway capacity and population abundance, rather than simply combining strains. The study’s contribution is the isolation and characterization of a fast paracetamol-transforming *Delftia* strain and the demonstration that its 4-AP accumulation can be reduced in a complementary co-culture, including in biologically treated wastewater. This strategy may help move wastewater treatment toward detoxification by reducing both the parent pharmaceutical and the persistence of an intermediate of concern. However, disappearance of paracetamol and lack of detectable 4-AP do not alone prove detoxification or mineralization. Future assessment should combine metabolite and carbon balances with toxicity measurements and should verify the proposed *sadB*-associated activity in the *M. lacus* strain. Within these limits, the findings show how literature-guided selection of complementary bacteria can help overcome a metabolic bottleneck and improve the prospects for biological removal of pharmaceutical contaminants.

## Supporting information

All the supplmentary materials supporting the article are included in the file.

## Acknowledgements

Acknoweldgement

I thank Vitali Helzer from FHNW, ICB institute for the SEM data.

## Availability Statement

The degradation data supporting the findings of this study are publicly available in Zenodo at https://doi.org/10.5281/zenodo.23922225.

## Author Contribution

F.D. planned the experimental setup, supported with lab work occasionally, performed the data analysis and supported the use of statistical models, drafted and revised the manuscript. M.SC. performed all experiments in the lab. C.Z. contributed on the isolation of *Delftia* P1. P.J., drafted and revised the manuscript. M.SU., and P.C revised the manuscript. All authors confirmed the final version of the manuscript.

## Declaration

## Ethical approval

This article does not contain any studies with humanparticipants or animals performed by any of the authors.

